# Machine learning of enhancer-promoter specificity based on enhancer perturbation studies reveals a distinct class of enhancers

**DOI:** 10.1101/2023.06.30.547290

**Authors:** Dylan Barth, Jonathan Cardwell, Mira V. Han

## Abstract

**Motivation:** Understanding the rules that govern enhancer-driven transcription remains a central unsolved problem in genomics. Now with multiple massively parallel enhancer perturbation assays published, there are enough data that we can utilize to learn to predict enhancer promoter relationships in a data driven manner.

**Results:** We applied machine learning to one of the largest enhancer perturbation studies integrated with transcription factor and histone modification ChIP-seq. Based on the learned model, we confirmed previously reported rules governing enhancer driven transcription, and we gained some insights that generated new hypotheses, such as a novel role for protecting against replication-transcription conflict at the active enhancers in CHAMP1. We also identified a distinct class of enhancers that drives target promoter transcription, but is not in strong contact with the promoters. There were two clusters of such enhancers that regulated *ATG2A* and the histone 1 cluster respectively. These enhancers were different from other typical enhancers, in that they had other strong enhancers nearby, and they also had strong H3K4me3 marks at the target promoters, both patterns that typically predict reduced enhancer influence, but here contributing in the opposite way. In summary, we find that integrating genomic assays with enhancer perturbation studies increases the accuracy of the model, and provides novel insights into the understanding of enhancer driven transcription.

**Availability:** the trained models and the source code are available at https://github.com/HanLabUNLV/abic.

## 1 Introduction

Enhancers are regulatory elements that affect gene expression through the direct physical interaction with promoters. This process often involves gathering various context specific transcription factors at the enhancer and the promoter where the general transcription factors bind to activate transcription. This general model has been validated through many studies (Palstra et al., 2003; Spilianakis & Flavell, 2004; Tolhuis et al., 2002), but it has also been challenged in certain contexts (Alexander et al., 2019). But, the majority of chromatin interactions between accessible regions does not involve a promoter. Instead, genes are surrounded by complex chromatin interactions mostly between enhancers creating a dense Chromatin Interaction Network (ChIN) (Miele & Dekker, 2008; Sandhu et al., 2012; Thibodeau et al., 2017). This framework has opened up the possibility of understanding how enhancers work cooperatively and indirectly in a network to fine-tune the level and timing of transcription (Lim & Levine, 2021; Rowley & Corces, 2018; Xiao et al., 2021).

A central question surrounding the interaction between regulatory elements and their target genes, is what principles determine which elements regulate any given gene and how such specificity is achieved. One working model that aims to solve this question is the Activity-by-Contact (ABC) model developed by Fulco et al. (Fulco et al., 2019). Based on solid biological principles, and a simple additivity assumption, the ABC model only requires 3 genome-wide measurements, H3K27ac, DHS, and Hi-C. The model allows the ranking of the enhancer’s importance to a gene’s regulation as proportional to its activity multiplied by contact. Fulco et al. performed CRISPRi flowFISH experiments and showed the utility of the model by predicting enhancer-promoter (EP) pairs with state-of-the-art accuracy (Fulco et al., 2019). While the ABC model is elegant, and works better than any alternative approach, it does not utilize the full availability of genomic data. Several important biological factors are ignored in the ABC model, such as the combination of transcription factors, the various chromatin marks other than H3K27ac surrounding the enhancer/promoter or the network of interactions in chromatin contact.

A 2019 paper by Song et al. focused on the hierarchical network of chromatin interactions and emphasized the role of what they called primary enhancers which came into direct contact with the promoter (Song et al., 2019). Song et al. used Hi-C data to create networks with edges representing contact between enhancers and promoters. Enhancers were then given labels based on how closely they were connected to a promoter on the network, such that *e_1_* was a step-one enhancer, directly in contact with the promoter, and *e_n_* was a step-*n* enhancer, n-1 steps removed. *e_1_*s were the main links between the promoter and other enhancers, evidenced by the fact that *e_1_*s had a higher betweenness centrality score than the rest of the enhancers in the chain (Song et al., 2019). Although showing a distinct pattern of contact between different enhancers, this study was limited by the fact that the enhancers they were studying were not functionally verified. This has been a difficult impasse for many studies interested in how enhancers regulate transcription on a large scale, but with the invention of CRISPR interference (CRISPRi), large parallel assays testing the functionality of individual enhancer elements are now possible.

Since Fulco et al. (Fulco et al., 2019) there have been several large scale studies that deploy CRISPRi to perturb multiple enhancers and measure the effects on gene expression often through single cell RNA-seq (Gasperini et al., 2019; Schraivogel et al., 2020). The accumulation of such data provides an opportunity to test the enhancer network model on verified enhancer data, and gain a more detailed understanding of different classes of enhancers. Here, we utilized the largest CRISPRi experiment to date (Gasperini et al., 2019) to predict functional enhancer-promoter pairs that showed significant downregulation after perturbation. Note that this aim is different from many studies that aimed to predict enhancer elements without relations to its target (de Almeida et al., 2022; Erwin et al., 2014; Yang et al., 2017). It is also different from numerous studies that came before high-throughput perturbation experiments such as TargetFinder (Whalen et al., 2016) or JEME (Q. Cao et al., 2017) that aimed to predict chromatin interactions (chromatin loops) without distinguishing functional effects on gene expression (F. Cao et al., 2021; Fudenberg et al., 2020; Schwessinger et al., 2020). The most relevant studies would be the recent study by Bergman et al. (Bergman et al., 2022) and Martinez-Ara et al. (Martinez-Ara et al., 2022) that systematically examined the compatibility of enhancers and promoters in human and mice respectively. But, the above studies utilized data from episomal high-throughput reporter assays that are different from CRISPRi experiments that occur in native genomic context. By training a machine learning algorithm on the CRISPRi experiments of Gasperini et al. (Gasperini et al., 2019), we could integrate various genomic assays measured on the candidate enhancer and target promoter in native genomic context into the feature space, such as the 3D chromatin contact and various TF ChIP-seq. Based on this framework, we aimed to predict functional enhancer-promoter relationships and learn the important features that contribute to enhancer-promoter specificity, especially in cases of direct and indirect contact.

## 2 Methods

### 2.1 Candidate Enhancer-promoter pair identification

To identify the potential candidate enhancer-promoter pairs in regulatory relationships, we decided to utilize the ABC software that will identify all enhancer and target promoter pairs within a 5Mbp window for every gene in the genome. Briefly, the ABC software defines a list of 500 bp candidate enhancers centered around DHS peaks, and filters out enhancers overlapping an annotated Transcription Start Sites (TSS) as promoters. We consider the terms TSS and promoter as interchangeable throughout this study. For candidate enhancers that will be mapped to Gasperini2019, we used the DNase data ENCFF001UWQ and ENCFF000SVO from ENCODE following (Gasperini et al., 2019). For candidate enhancers that will be mapped to Fulco2019, we used the DNase data wgEncodeUwDnaseK562AlnRep1 and wgEncodeUwDnaseK562AlnRep2 from the UCSC genome browser.

### 2.2 Chromatin Interaction Network (ChIN) generation

We generated a chromatin interaction network based on Hi-C contact maps centered around the promoter for each gene, where each node was an enhancer or promoter identified by the ABC software above, and the edge weights of the network were equal to the KR-normalized Hi-C contact between those nodes. At this stage, each gene has a very dense network that is both computationally inefficient to traverse and expensive to store in memory (Fig. 1A). For this reason, we decided to simplify the network with edges that correspond to detectable chromatin loops. We used FitHiC2 (Kaul et al., 2020) with a cutoff of FDR<0.01 to call significant loops, and only kept the contact edges corresponding to loops, filtering out all edges without a significant loop detected (Fig. 1B). Once these simpler networks were created, they were further filtered by only keeping edges connecting enhancers and the promoters that were tested in Gasperini et al. (Gasperini et al., 2019). From these simplified networks, the degree of each enhancer with respect to each nearby promoter was determined following (Song et al., 2019). In short, enhancers that were directly connected to the promoter with a significant Hi-C loop were labeled *e_1_*s, and enhancers that were not connected to the promoter, but were connected to an *e_1_* were labeled *e_2_* and so on. Enhancers within the same Hi-C window as the promoters were labeled *e_0_*s, and enhancers that were more than 3 degrees away from the promoter or were not part of that promoter’s subcomponent of the graph were labeled *e_inf_*.

**Figure 1.**
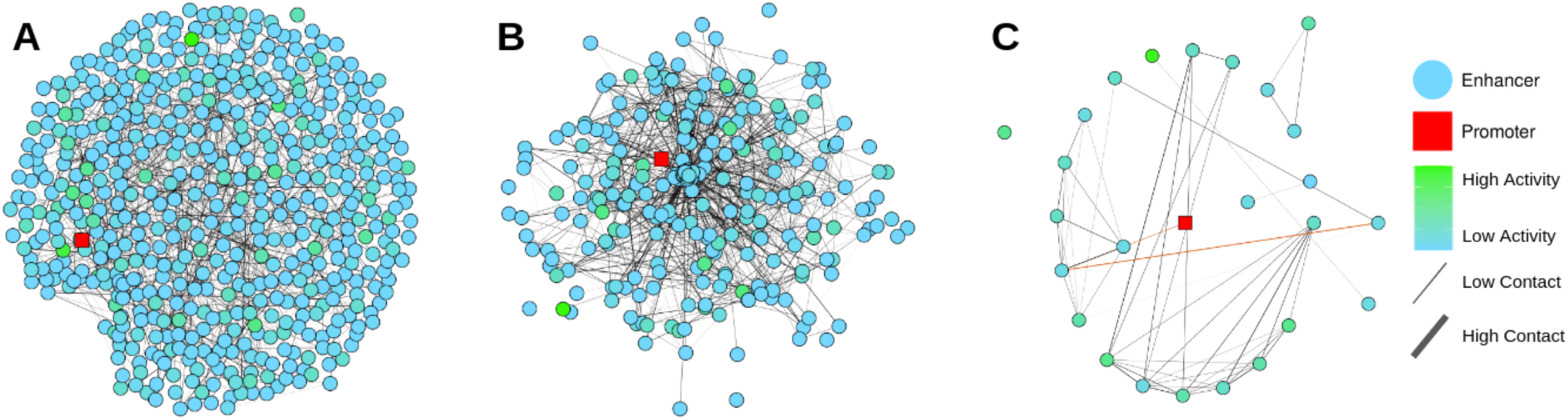
Chromatin Interaction Network (ChIN) generation process. Chromatin Interaction Network generation process of the HIST1H2BD gene and its highly dense surrounding Hi-C contact network. Nodes are candidate enhancers identified based on open chromatin, and edges are the KR-normalized Hi-C contact scores between two nodes. The promoter is shown in red and enhancer nodes are color scaled based on their ABC activity score (green; high, blue; low). **A)** Unfiltered network showing all Hi-C contacts. **B)** Network with edges filtered based on significant Hi-C contact (FitHiC2 q-value < 0.01). **C)** Network with nodes filtered based on experimental data. Only experimentally tested (Gasperini et al., 2019) enhancer nodes are present in graph. The path connecting a significant *e_2_* enhancer to the promoter is highlighted in red.

### 2.3 Feature Engineering

The input to the machine learning pipeline consists of a feature matrix whose rows are enhancer-promoter pairs, and whose columns represent the variables measured for each entry. The features (variables) include measures of histone modification (H3K27ac, H3K4me3, H3K27me3) around the enhancer, the same histone modifications around the TSS, expression level of the target gene, genomic distance between the enhancer and the TSS, Hi-C contact measure between the enhancer and the TSS, the denominator - numerator of the ABC score that we call the rest-of-ABC representing strengths of other enhancers nearby, the various TF presence at the enhancer measured from 250 ChIP-seq experiments, the 250 TF presence at the TSS, and whether a same TF is bound at the enhancer and the TSS.

The histone modification peaks were obtained from the histone modification ChIP-seq data from ENCODE (ENCFF384ZZM for H3K27ac, ENCFF681JQI for H3K4me3, ENCFF937NEW for H3K27me3) and processed following the approach of Fulco et al. The enhancer activity is quantified based on a combination (geometric mean) of DNase-seq and H3K27ac ChIP–seq signals following the approach of Fulco et al.

The Hi-C scores were obtained from the .hic matrix provided by Aiden et al. (https://hicfiles.s3.amazonaws.com/hiseq/k562/in-situ/combined_30.hic) and processed following the approach of Fulco et al. Hi-C contact scores were calculated based on the Hi-C signal between the enhancer and the TSS of the gene. We note here that although we relied on the ABC software for the candidate enhancer list and the normalization of histone modification marks and Hi-C intensities, we didn’t use the final ABC score in our analysis, because we were interested in how individual features contributed to the prediction.

The transcription factor peaks were obtained from the “Transcription Factor ChIP-seq Clusters from ENCODE 3” track from the ucsc genome browser, summarized in a downloadable bed format at https://hgdownload-test.gi.ucsc.edu/goldenPath/hg19/encRegTfbsClustered/encRegTfbsClusteredWithCells.hg19.bed.gz. We extracted the peaks for the K562 cell type from the file and used bedtools to intersect the peaks of each TF with the set of all enhancers and promoters. The peaks were encoded into integers indicating the number of peaks present within the 500bp region. This process culminated in a set of 768 features recorded for 66,283 enhancer-promoter pairs based on a set of 250 TFs. The full list of features can be found in Supplementary table 1.

### 2.4 Training data, validation data and Independent test data

To compile experimentally verified positive and negative enhancer-TSS pairs that we used for training, validation and test, we overlapped our candidate enhancers and promoters with experimental data from two independent CRISPR perturbation studies, Gasperini2019 (Gasperini et al., 2019) and Fulco2019 (Fulco et al., 2019). The output we aimed to predict is a binary class label that was assigned based on the experiments in Gasperini2019 (or Fulco2019) that is positive (equal to 1) only if blocking said enhancer using a CRISPRi gRNA resulted in significant (FDR<0.1) down regulation of the gene. The parameters for the bedtools intersect command (*bedtools intersect -wo -e -f 0.3 -F 0.3 -a*) were determined based on the fraction parameter that gave us the maximum intersection between our candidate enhancers and Gasperini2019. Within Gasperini2019, some of the enhancers were split into smaller segments targeted by different guide RNA, so there were a few *n:*1 relationships between the *n* smaller Gasperini2019 enhancers all contained within a single larger (500bp) candidate enhancer extracted from DHS peaks. For these, if any of the *n* enhancers were positive in the experiment, the whole candidate enhancer was called positive in our compiled data. Within Fulco2019, a few of the enhancers were combined and merged in their final table, leading to longer than expected enhancers. This resulted in both *n*:1 and 1:*n* relationships between enhancers tested in the experiment, and our candidate enhancers. In case of *n*:1, we followed the same rule, and considered our candidate enhancer positive if any of the contained enhancers tested were positive. In case of 1:*n*, all of our candidate enhancers were retained in our dataset, but they were all assigned the same positive or negative outcome based on the Fulco2019 results.

The compiled dataset of Gasperini2019 and the associated features was divided into training and test data, by setting aside chromosomes 5, 10, 15, and 20 as test sets. An independent second set of test data was generated by compiling the Fulco2019 and the associated features in the same manner. The remaining training data of Gasperini2019 (chromosomes other than 5, 10, 15, 20) was then divided into a 4 x 4 nested blocked cross validation scheme (using *StratifiedGroupKFold*), where blocking was based on the groups defined by at least 5Mb of gap separating any of the enhancers or TSS in the data. Any enhancer-TSS pairs that are within the block were assigned to the same group, and all data within the same group were assigned into a single fold, ensuring separation by genomic distance, and thus independence between data across each fold (Whalen et al., 2022; Xi & Beer, 2018). The grouping of the data and folds can be found in Supplementary Table 2. The outer folds were used for evaluation of performance, while the inner folds were used for hyperparameter optimization.

### 2.5 Machine learning pipeline

We used gradient boosting (XGBoost) for the learning algorithm (Chen & Guestrin, 2016). We explored other algorithms that didn’t give us similar or better performance, so we don’t report those results. As described above, the outer folds were used for evaluation of performance, so all of the scaling, feature selection, hyper-parameter optimization was done within the training split of the outer fold. Test split of the outer fold was used for unseen performance evaluation for the pipeline trained on the train split. We didn’t retrain the final pipeline on combined folds. We opted to evaluate the 4 different pipelines produced by 4 folds separately, to get a measure of uncertainty around our performance evaluation.

Within each train split, we scaled the features using MinMaxScaler to be between 0 and 1. We did feature selection using Boruta with SHAP values as importance measures on gradient boosted trees trained through a preliminary optimization (Kursa & Rudnicki, 2010). We were more lenient in retaining the features by adjusting the max shadow features importance measure by 80%, and using a p-value cutoff of 0.1. The hyperparameters of the XGBoost classifier were optimized on the inner CVs using Optuna (Akiba et al., 2019), with pruning and early stopping based on the validation set metric. Due to the extreme imbalance in our data (605/65,696 positive cases), we used Mean Average Precision (MAP) averaged across the inner folds as the metric to maximize during CV. We used the default algorithm of Tree-structured Parzen Estimator, and the MedianPruner on the validation MAP. The search space we explored and the final optimized parameters can be found in Supplementary Table 3.

The SHAP values were calculated using the tree SHAP implementation in XGBoost. SHAP values for the four test folds from the four models were combined through concatenation.

## 3 Results

### 3.1 Enhancer network characteristics

In total, we generated 7,396 gene networks. After filtering the edges by calling significant loops (FDR<0.01) using the FitHiC2 software (Fig. 1B), and pruning out any nodes that did not overlap with the Gasperini data (Fig. 1C), the networks had an average of 7.4 nodes and 55.8 edges (Fig. 2A, 2B). Most of these edges did not connect to the promoter, as on average the degree of the promoter is only 4.9. Despite having few edges directly connected to the promoter on average, the promoter node was often very close to the rest of the nodes in the network. Closeness centrality is a measure of how interconnected a node is within a network and is calculated as approximately the reciprocal of the shortest distance to every node in the network. Values near 1 indicate that the node is separated from the rest of the network by small distances on the graph, and values near 0 indicate that the node is separated from the network by large distances. The majority of the promoters in our networks tend toward a closeness measure of 1, underscoring their importance within the network despite a lack of direct connections to most nodes in the network (Fig. 2C).

**Figure 2.**
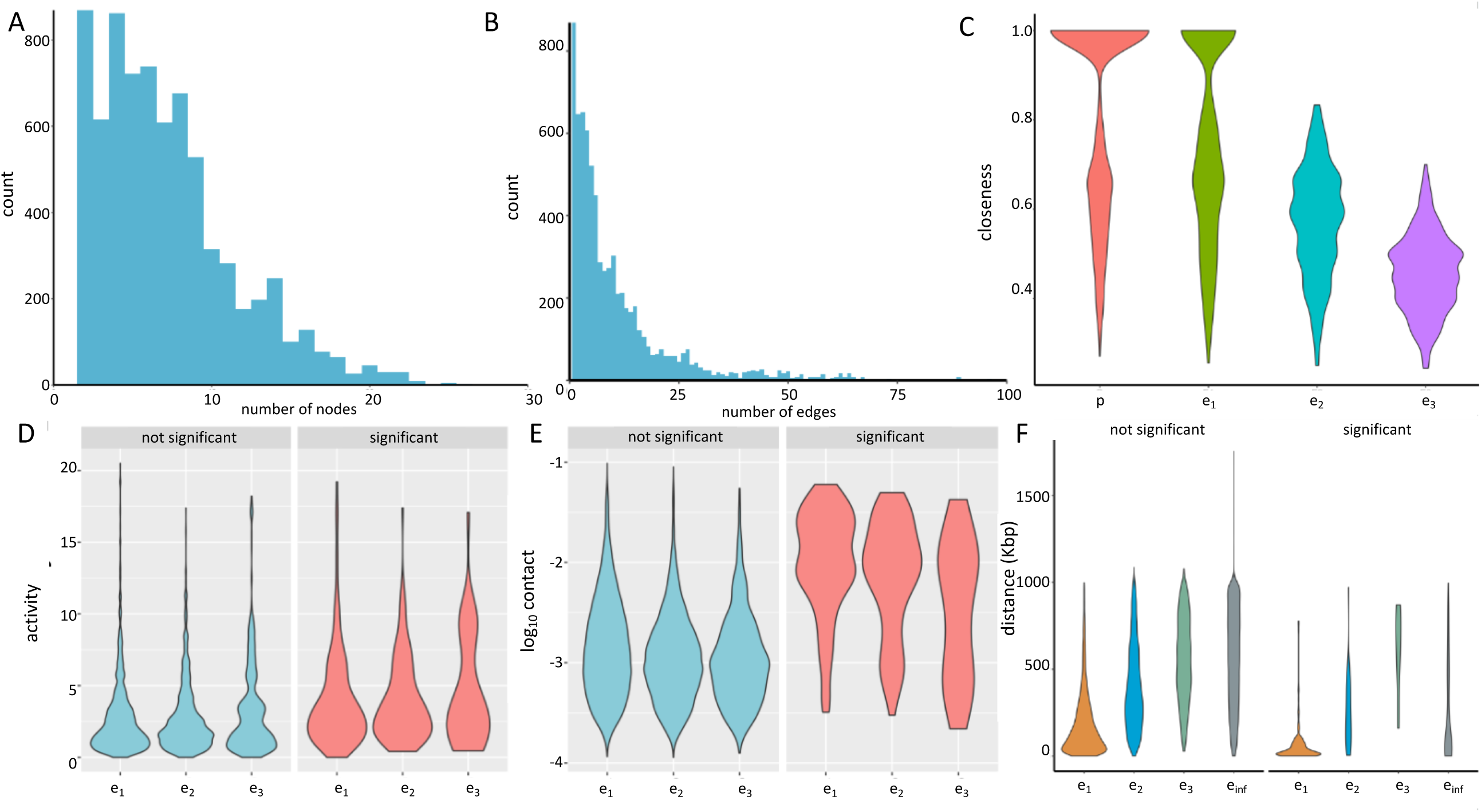
ChIN and enhancer characteristics. Histograms showing distribution of the number of nodes **(A)** and edges **(B)** for each network. **C)** Closeness measure of each class, *e_1_* and promoter show higher closeness centrality to each network. Activity **(D)** and Contact **(E)** within each class of enhancer delimited by functional significance in Gasperini et al. 2019. **F)** Linear distance between each class of enhancer and their corresponding promoters measured in kbp.

### 3.2 Enhancer characteristics

Out of 65,696 enhancer-promoter pairs that were tested by Gasperini2019, 605 were positive, *i.e.* perturbation of the enhancer was found to significantly alter the expression of the target gene. Of those 605 significant EP pairs, 26 were classified as *e_0_*, 347 were classified as *e_1_*, 29 were classified as *e_2_*, 11 were classified as *e_3_*, and the remaining 192 were classified as *e_inf_*, as they were either more than 3 steps away from the promoter on the network or they were not significantly connected to the promoter at all using our FitHiC2 threshold of FDR<0.01. An example of an *e_2_* is shown in Fig. 1C.

Despite the ABC model not taking into account these higher degrees of enhancer contact, the essence of the model can be seen by comparing the activity and contact of each enhancer class (Fig. 2D-E). As expected, the further away from the promoter on the network, the lower the contact with the promoter is. Focusing specifically on the significant EP pairs reveals that for an enhancer to remain functional despite having lower contact with the promoter, it is required to have a slightly higher activity level to compensate. Song et al. (Song et al., 2019) noticed a distinction that *e_2_* were much closer to the promoters in linear genomic distance compared to *e_1_*, which we did not observe (Fig. 2F).

### 3.3 Integration of additional genomic features improves prediction of functional enhancer-promoter pairs

We evaluated the performance of the trained algorithm on various test datasets to get a robust assessment. The overall mean average precision from the inner fold validations were around 0.26∼0.28, and the outer fold average precision on the test folds ranged from 0.21∼0.30 (Table 1A). Although the outer test fold performance has a larger variance, the overall performance between the CV and the test is comparable, showing that we are not overfitting to the training data. We also had an independent test set from Gasperini2019 consisting of chromosomes 5,10,15,20 that were set aside from the nested cross validation design, and the test performance on those data also showed comparable values of average precision between 0.23-0.29, again showing estimates of performance without overfitting (Table 1C).

**Table 1.**
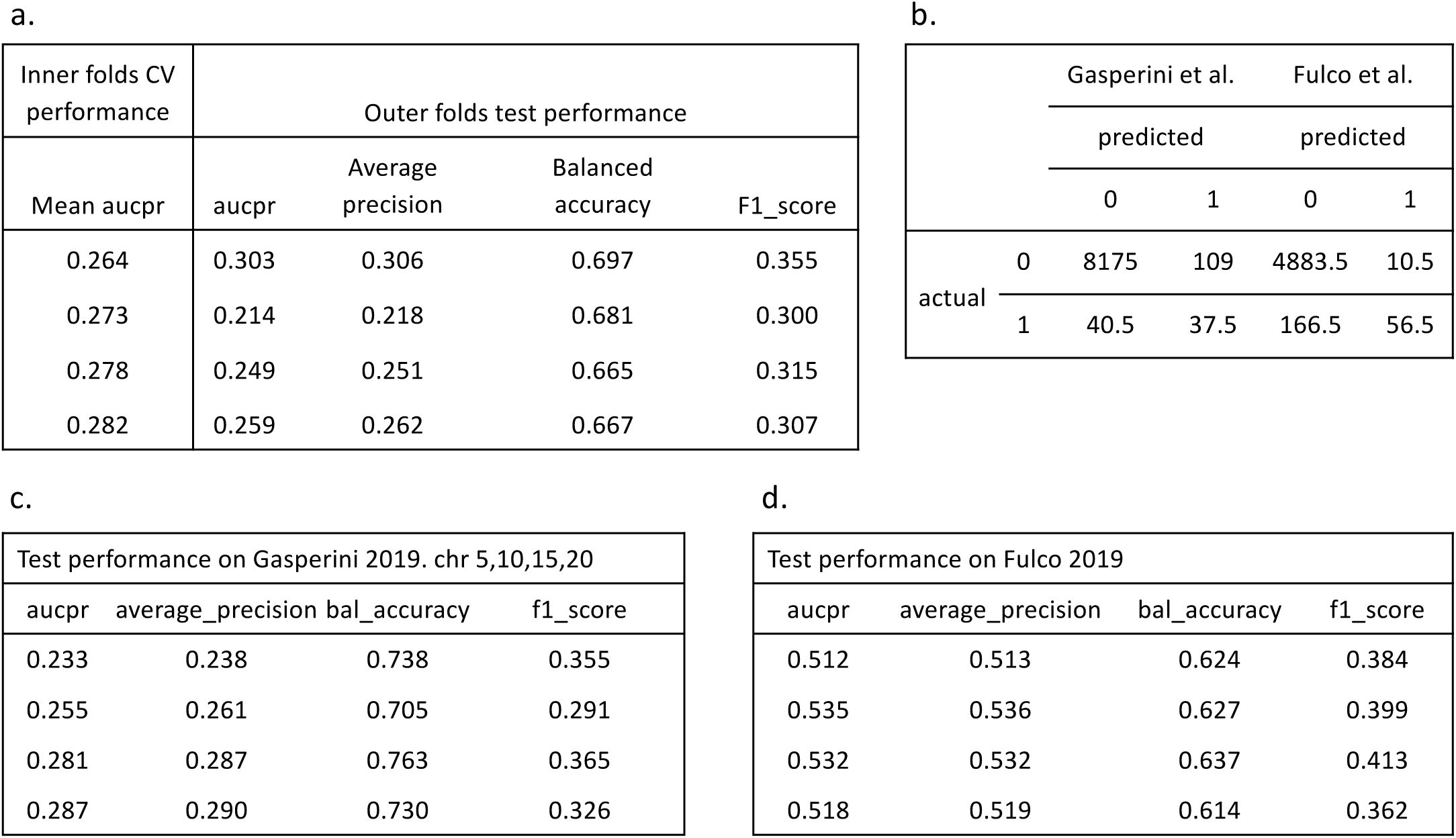
Model performance. Performance of learned model on Gasperini2019 and Fulco2019 datasets. **A)** Inner fold CV performance and outer fold test performance for each of the four models trained on the four folds. **B)** Confusion matrix produced by the model on each test data averaged across the four models. **C)** Test performance on the data from Gasperini et al. 2019. chromosomes 5,10,15,20. **D)** Test performance on the data from Fulco et al. 2019

We find that integrating various genomic measures as features in the learning increased the performance of enhancer-promoter prediction compared to the ABC model or distance-based prediction. On the Gasperini2019 dataset, we are achieving a modest Average Precision of around 0.269, with as many false negatives and more than twice false positives as the number of true positives (Table 1B). But considering the challenge of the dataset, which is an extremely imbalanced dataset with less than 1% positive rate (78 positive out of 8326 total in the chr 5,10,15,20 test data), we see a meaningful increase in the performance (Fig. 3A). With the Fulco 2019 dataset, which has a higher positive rate of about 4.3% (223 positive out of 5117 total), the performance reaches 0.51∼0.54 average precision (Table 1C). We see again a meaningful increase in the performance compared to ABC or distance (Fig. 3B).

**Figure 3.**
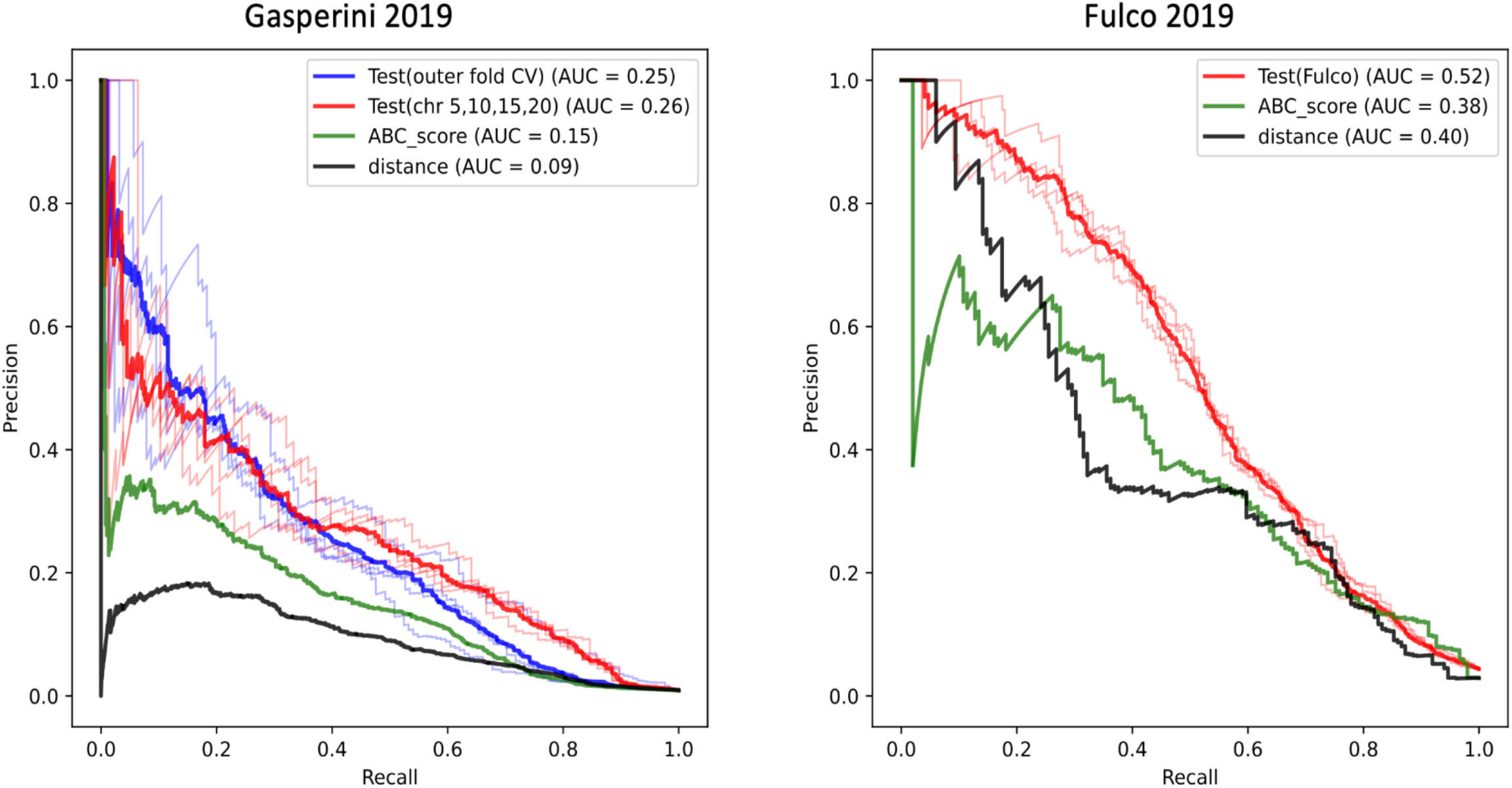
Comparison of model performance. Precision-recall curves of various models on **A)** Gasperini2019 and **B)** Fulco2019 datasets. In every case, our model trained on the integrated genomic data with the XGBoost algorithm outperforms both distance based and ABC predictions.

### 3.4 Feature importance of the learned model corroborates the ABC model and other rules of transcriptional regulation

To understand where the increased performance is coming from, and gain insight into the prediction of functional enhancer-promoter pairs, we tried to explain the trained model using Shapley values (SHAP) (Lundberg & Lee, 2017). The SHAP values show the features that are important in driving the model’s prediction of functional EP pairs. Positive SHAP value means positive impact on prediction, leading the model to predict 1 (*i.e.* perturbation of enhancer results in downregulation of target gene). Negative SHAP value means negative impact, leading the model to predict 0 (*i.e.* perturbation of enhancer did not have effect on target gene). Note that positive enhancer-promoter pairs will tend to have large negative effect size in their differential gene expression after perturbation. To avoid confusion between the terms, we will sometimes use the terms functional vs. non-functional EP pairs, instead of positive vs. negative EP pairs, when we are discussing effect size at the same time. To summarize the SHAP values that we obtained, stronger Hi-C contact, shorter distance between enhancer and target TSS, weaker neighboring enhancers near the TSS, higher target gene expression, higher H3K27ac peak at the enhancer, all contributed to the model to predict positive enhancer-promoter pairs (Fig. 4). The factors that are components for the Activity by Contact model such as the Hi-C contact, and the H3K27ac chromatin marks at the enhancer were at the top of the feature ranking, confirming the validity and power of the ABC model. In addition, the enhancer-promoter is more likely to be positive when the value of the rest-of-ABC score (ABC denominator – numerator) is lower, *i.e.* when the neighboring enhancers nearby are weaker (although we see some interesting exceptions that we describe below). This also justifies the approach of ABC, which compares the relative effect of each enhancer, normalized by the sum of ABC of other nearby elements.

**Figure 4.**
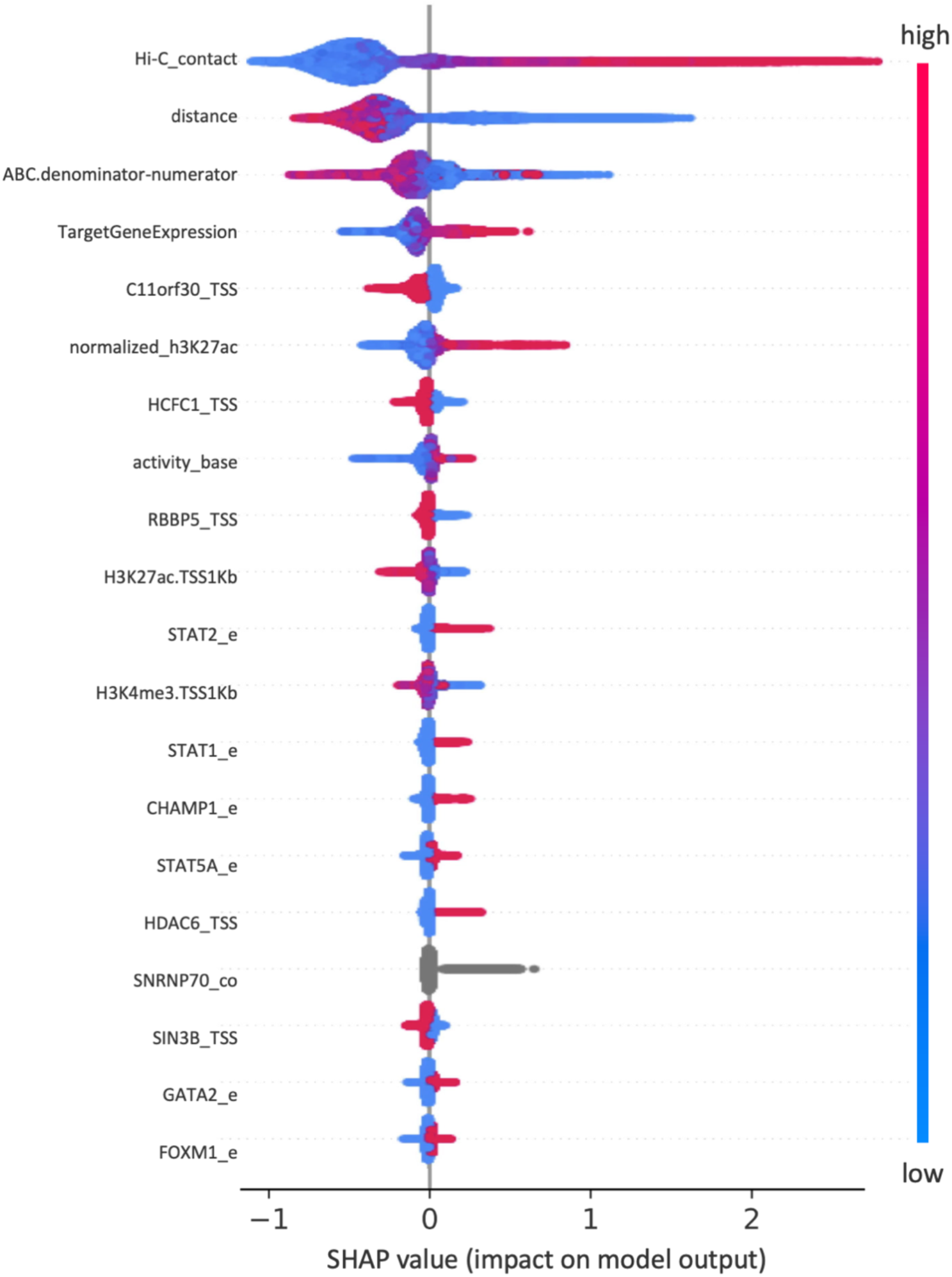
SHAP values. SHAP values of most important features ranked from top to bottom by their mean absolute SHAP values. Every sample of the data appears as a point along the horizontal axis. Colors correspond to the measured value of the input data for that variable. The X axis predicts the outcome as positive or negative. For example, high values of Hi-C contact (red) predicts positive outcome (right of the horizonal axis), while low values of distance (blue) predicts positive outcome (right of the horizontal axis). Here positive outcome means functional E-P pairs where CRISPRi has a downregulation effect.

In addition to explaining important features, the boosted trees combined with SHAP analysis gave us insight into the quantitative threshold values of the features that lead to different predictions. For example, the dependence plot of Hi-C contact and H3K27ac shows that Hi-C contact, the most important feature, doesn’t need to be very large in value. The SHAP value rises with a steep slope. Even with a KR-normalized Hi-C contact frequency as small as 0.01, the pairs are more likely to be positive, and the SHAP values plateau by the Hi-C value of 0.03 (Fig. 5A-B).

**Figure 5.**
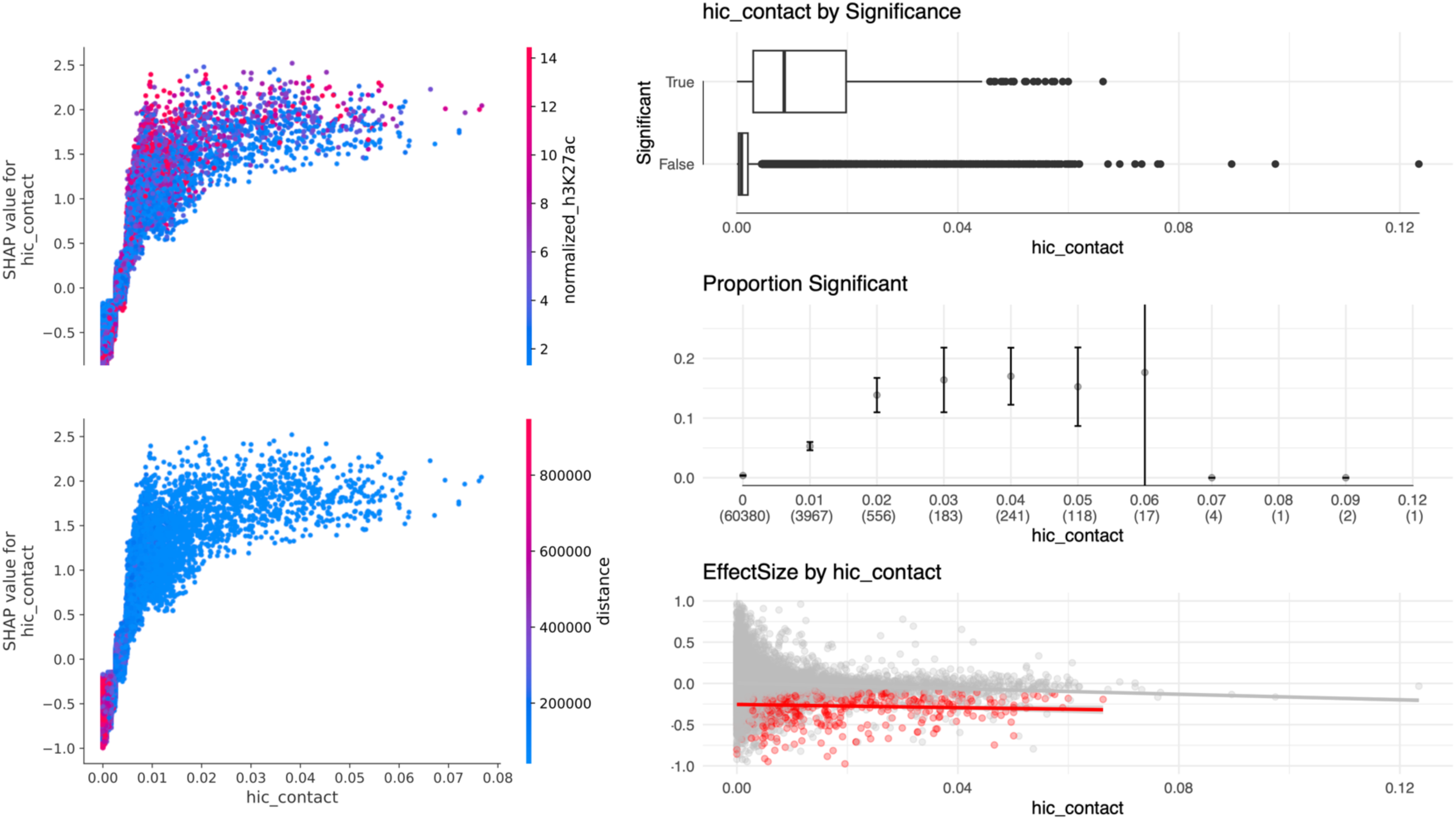
Strong Hi-C contact predicts functional enhancer-promoter pairs. **A)** Hi-C contact shows positive trend with SHAP values, and interaction with H3K27ac at the enhancer. Vertical pattern of coloring for the same values of Hi-C contact reveals interaction between Hi-C and H3K27ac, recapturing the dynamics of the ABC model. **B)** SHAP values for Hi-C ontact colored by genomic distance between enhancer and promoter. **C)** Contact by significance boxplot which shows functional enhancers are in strong Hi-C contact with their corresponding promoters more often than nonfunctional enhancers. **D)** The proportion of significant data increases as Hi-C contact increases. The highest proportion of significant data in our dataset has a normalized Hi-C score of 0.06. Numbers in parentheses show the number of samples (n) in each Hi-C bin. **E)** Effect size vs Hi-C score (significant samples in red). Significant enhancer-promoter pairs with stronger Hi-C contact have larger negative effect size (larger down-regulation from CRISPRi on paired enhancers).

Since SHAP values are based on the trained predictive model, we explored whether the signal can be seen in the actual data itself. Data from Gasperini2019 and Fulco2019 both showed large differences in the Hi-C contact strengths between positive pairs and negative pairs (Fig. 5C). As seen from SHAP values we saw an increase in the proportion of significant pairs as Hi-C value increases to as low as 0.01 and we observe the plateau by Hi-C contact value of 0.03. (Fig. 5D). Among the functional EP pairs, the effect size shows larger negative values with stronger Hi-C contact (Fig. 5E). Here, larger negative effect size means stronger down-regulation with CRISPR interference, *i.e.* stronger functional effect.

Another interesting feature highlighted by the SHAP analysis is the H3K27ac and H3K4me3 chromatin marks at the TSS. Both chromatin marks contribute to negative prediction, meaning that if there is a strong activating mark at the TSS, the model is less likely to predict a functional enhancer-promoter pair for that TSS (Fig. 6A-B). Initially this seems unexpected, but it is consistent with several observations that have been reported on transcriptional regulation. Bergman et al. showed that promoters of ubiquitously expressed (housekeeping) genes, which they called P2 promoters, have higher H3K27ac marks at the promoter and are less responsive to distal enhancers (Bergman et al., 2022). Similarly, in both Gasperini2019, and Fulco2019, the experiments showed that housekeeping genes are depleted from their positive enhancer-promoter pairs, indicating that housekeeping genes are less influenced by enhancers, and their transcription is sufficiently driven by promoters alone.

**Figure 6.**
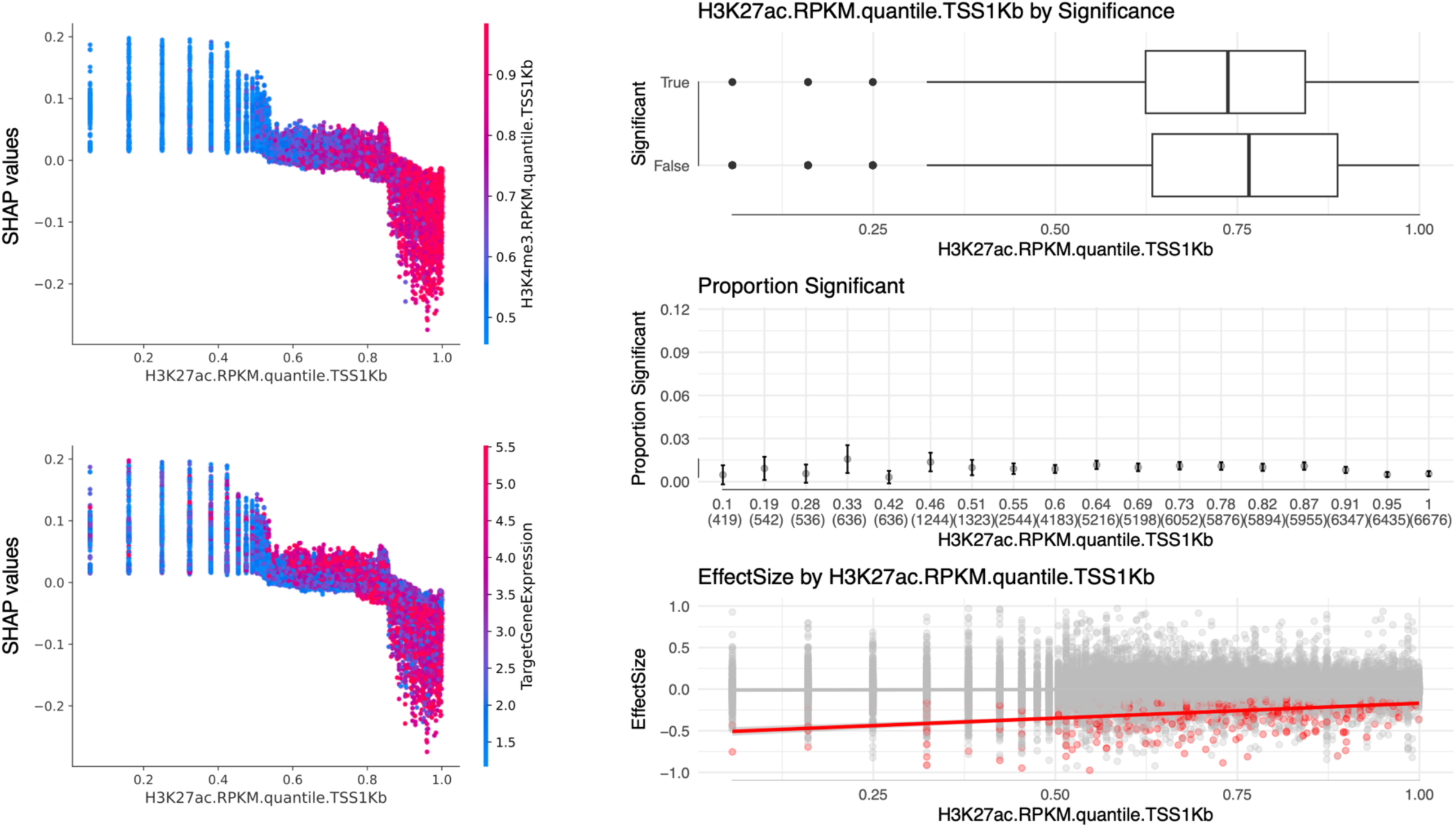
Strong H3K27ac at the promoter predicts nonfunctional enhancer-promoter pairs. Promoter H3K27ac mark shows negative trend of SHAP values. Promoter H3K27ac shows correlation with promoter H3K4me3 **(A)** and target gene expression **(B) C)** Promoter H3K27ac by significance boxplot. Significant enhancer-promoter pairs show decreased promoter H3K27ac. **D)** There is reduced significance among the target genes with the strongest H3K27ac marks at promoters. **E)** Effect size vs H3K27ac strength (significant in red). Strong H3K27ac promoters have smaller negative effect size (weaker down-regulation effects from CRISPRi on paired enhancers).

### 3.5 Transcription factors present at the enhancers predict functional enhancer-promoter pairs while EMSY and HCFC1 present at the TSS predict nonfunctional enhancer-promoter pairs

The transcription factor ChIP-seq peak presence at the TSS and enhancers were not as important as the Hi-C interactions or chromatin marks at the TSS and enhancers, but they contributed to further increase the predictive performance. The most important TFs at the TSS and enhancers contributed in different directions, where the TFs present at the TSS predictions negative (nonfunctional) EP pairs, and TFs present at the enhancers predicted positive (functional) EP pairs. The two most important TFs are C11orf30 (EMSY) at the TSS, and HCFC1 (HCF1) at the TSS, which were both associated with nonfunctional enhancer-promoter prediction (Fig. 7, 8). Since EMSY and HCFC1 each interact with the *Sin3* histone deacetylase (HDAC) respectively, and EMSY is known as a repressor, we initially thought this is through the repressive action of EMSY and HCFC1 at the promoter. But, unexpectedly, Both EMSY and HCFC1 peak at the TSS were associated with higher H3K27ac marks and H3K4me3 marks at the TSS, and more active promoters (Fig. 7).

**Figure 7.**
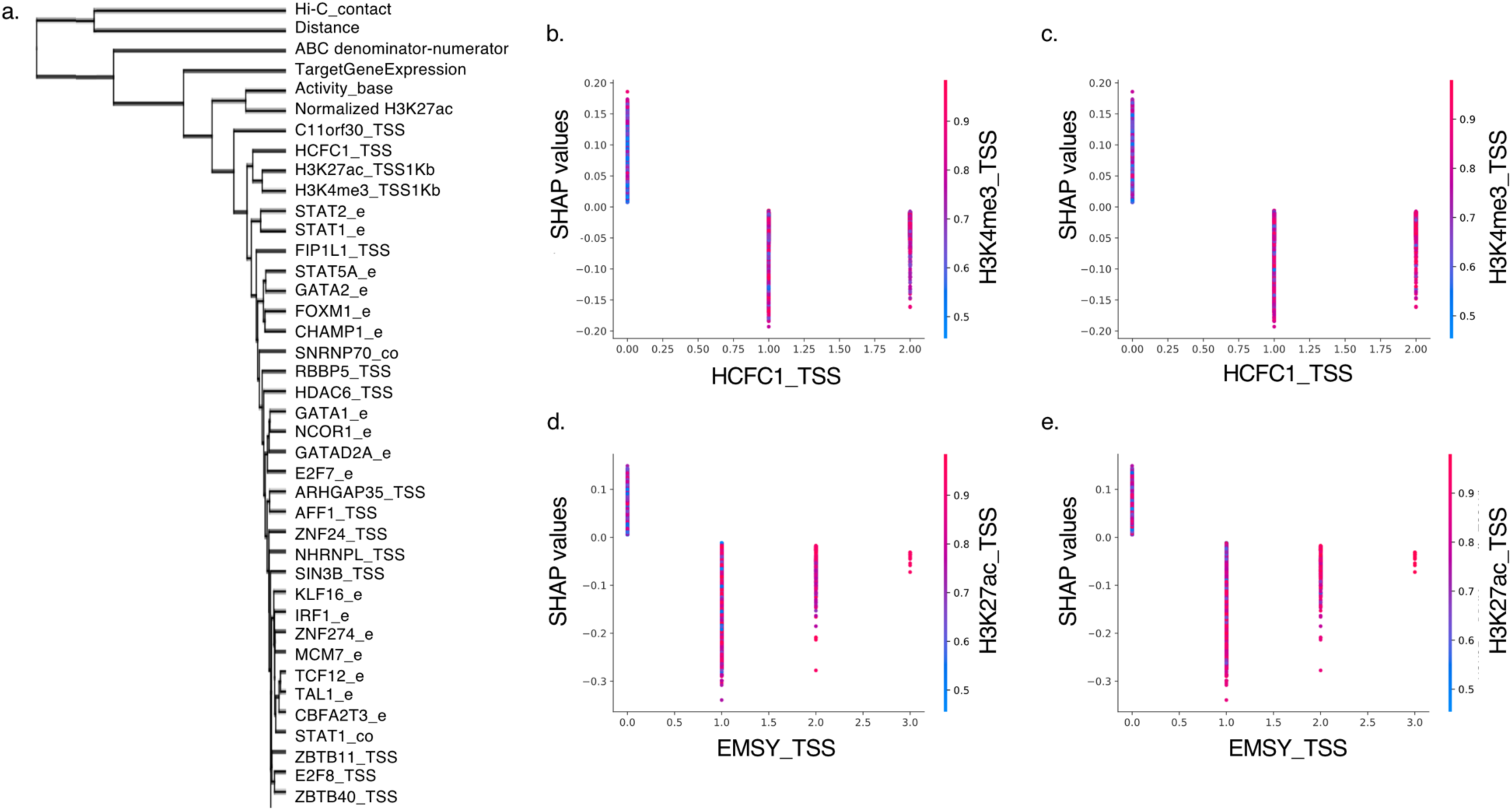
EMSY(C11orf30) and HCFC1 at the promoter predicts nonfunctional enhancer-promoter pairs. **A)** Hierarchical clustering of SHAP features based on their contribution to prediction. HCFC1 at the TSS is correlated with H3K27ac and H3K4me3 at the TSS in predicting nonfunctional EP pairs. **B-E)** SHAP values of *HCFC1* and *EMSY* (*C11orf30*) at the promoters. Presence of *HCFC1* and *C11orf30* peaks show correlation with both H3K27ac and H3K4me3 marks at the promoters, and show lower SHAP values indicating prediction of nonfunctional EP pairs.

This was also confirmed through the hierarchical clustering of SHAP values across feature variables. The clustering shows that H3K27ac, H3K4me3, and HCFC1 at the TSS are correlated in their effect on prediction across features and samples (Fig. 7A). The correlation between HCFC1 and to a lesser extent EMSY with both chromatin marks at the TSS are also found in the ChIP-seq and chromatin data that shows higher (quantile normalized) H3K27ac and H3K4me3 chromatin marks when either of the TFs are present. As expected, the ChIP-seq peaks between the two TFs are also correlated, co-occurring more often than by chance. Since SHAP values reflect correlation, not causation, we cannot disentangle the causal relationship between these four variables. But, we can tell that there are correlation among H3K27ac, H3K4me3, HCFC1 and EMSY peaks at the TSS that all eventually lead to decreased prediction of functional enhancers affecting the target gene.

On the other hand, numerous transcription factors found at the enhancer region were predictive of the functional enhancer-promoter pairs in the positive direction. The most important TFs found are the well-known hematopoietic and/or cancer related TFs, STAT2, STAT1, CHAMP1, STAT5A, GATA2, FOXM1, etc. (Fig. 4, Fig. 9)

**Figure 8.**
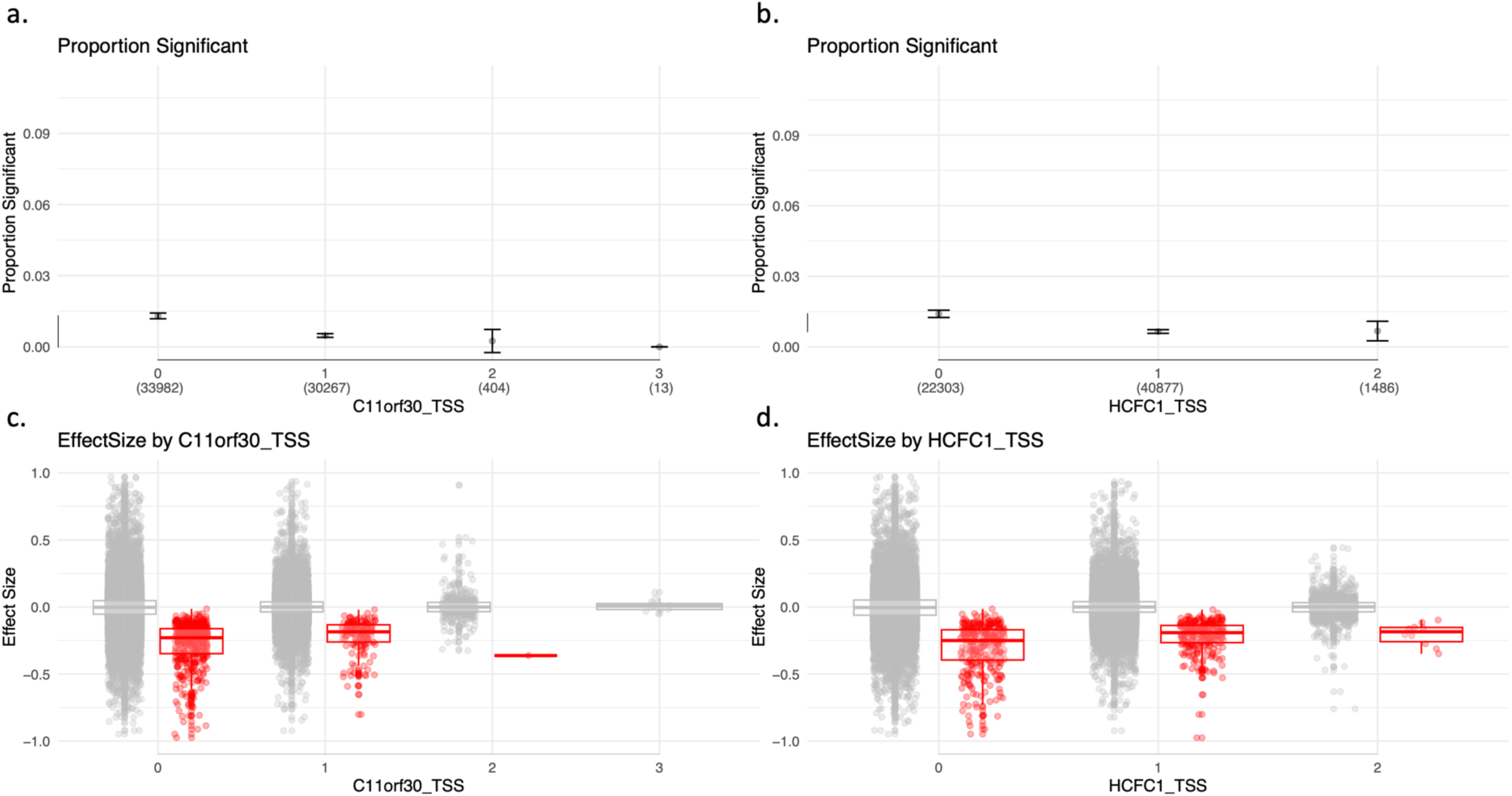
EMSY(C11orf30) and HCFC1 at the promoter is associated with less significance and smaller effect size. **A)** Presence of ChIP-seq peaks of EMSY (*C11orf30*) and **B)** *HCFC1* found at promoters are associated with lower proportion of significant enhancer-promoter pairs. **C)** Effect sizes of data stratified by the number of *C11orf30* and **D)** *HCFC1* at promoter peak. The presence of EMSY or HCFC1 at the promoter are associated with smaller negative effect size (weaker down-regulation effects from CRISPRi on paired enhancers).

**Figure 9.**
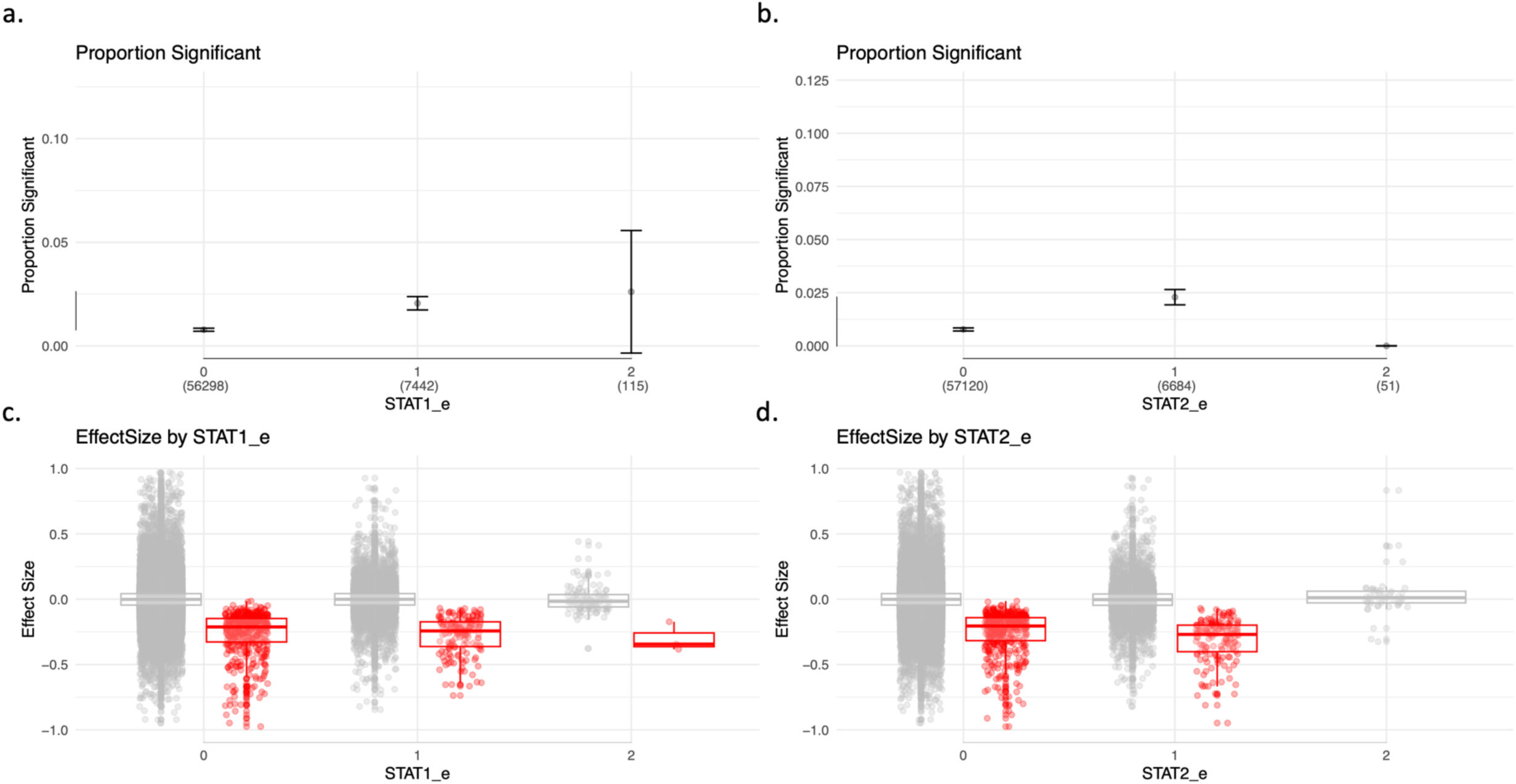
STAT1 and STAT2 at the enhancer predicts functional enhancer-promoter pairs. **A)** presence of ChIP-seq peaks of *STAT1* and **B)** *STAT2* found at enhancers are associated with higher proportion of significant enhancer-promoter pairs. Effect sizes of EP pairs stratified by the number of **C)** *STAT1* and **D)** *STAT2* peaks at the enhancer. The presence of STAT1 or STAT2 at the enhancers are associated with larger negative effect size (stronger down-regulation effects from CRISPRi on the enhancers).

The only gene that shows up as important for prediction that is relatively less studied in the context of hematopoiesis or cancer would be CHAMP1, a gene associated with a neurodevelopmental disorder (Hempel et al., 2015; Itoh et al., 2011). CHAMP1 interacts with Rev7, HP1, and POGZ and functions in chromosome segregation and DNA repair (Li et al., 2022). We found that CHAMP1 presence at the enhancer is correlated with FOXM1 presence at the enhancer (correlation coefficient 0.428) and they both increase the prediction of positive enhancer-promoter pairs. Given the function of the gene, it is possible that the gene is involved in protecting the DNA during replication transcription conflict, and functional enhancers are especially at risk for replication stress in this cancerous system.

### 3.6 Sorting the enhancers based on chromatin interaction networks reveal a distinct class of enhancers

Machine learning revealed that chromatin interaction measured by Hi-C contact between the enhancer and the promoter is the single most important feature in predicting functional enhancer-promoter pairs. Nevertheless, there are a substantial number of positive functional pairs that are not in strong contact. Gasperini et al. discussed this observation as underscoring the difficulty of the prediction task. In order to understand the difference between functional pairs that are in strong chromatin contact, and functional pairs that are not in contact, we split the data into two classes, direct contact enhancers that we call e1minus (because it encompasses *e_1_* enhancers that are in significant contact with TSS based on fitHiC2 calls, and *e_0_* enhancers that are within the same 5Kb window of Hi-C), and indirect enhancers that we call e2plus (*e_2_*, *e_3_*, *e_inf_*), that are not in direct contact with the TSS in the chromatin interaction network. We trained the same machine learning pipeline on these two subsets of data. Overall, we found that limiting the data to e1minus enhancers with direct contacts (341/10663=3.2% positive rate) made the prediction easier, reaching average precision that is about 30% higher than what we saw with the whole data (Table 2A). In contrast, the training and prediction of e2plus enhancers without direct contact (167/43102=0.4% positive rate) were extremely challenging, resulting in overfitting based on the difference seen between Cross-Validation MAP and Test MAP, and an order of magnitude lower average precision (Table 2B).

**Table 2.** Model performance for model trained on EPs with vs without direct Hi-C contact. performance measure by Average Precision for model trained on **A)** EPs with direct contact (e1minus data) and **B)** EPs without direct contact (e2plus data).

| Inner folds<br>CV<br>performance |  | Outer folds test performance |  |  |
| --- | --- | --- | --- | --- |
| Mean aucpr | aucpr | average_<br>precision | balanced_<br>accuracy | f1_score |
| 0.363 | 0.325 | 0.330 | 0.627 | 0.335 |
| 0.368 | 0.226 | 0.230 | 0.595 | 0.249 |
| 0.349 | 0.466 | 0.469 | 0.729 | 0.462 |
| 0.376 | 0.276 | 0.290 | 0.669 | 0.350 |

| Inner folds<br>CV<br>performance |  | Outer folds test performance |  |  |
| --- | --- | --- | --- | --- |
| Mean aucpr | aucpr | average_<br>precision | balanced_<br>accuracy | f1_score |
| 0.088 | 0.029 | 0.033 | 0.521 | 0.063 |
| 0.125 | 0.064 | 0.078 | 0.562 | 0.151 |
| 0.085 | 0.059 | 0.038 | 0.531 | 0.061 |
| 0.085 | 0.039 | 0.036 | 0.532 | 0.079 |

Despite the lower performance of e2plus enhancers, the overall feature importance based on SHAP values showed generally similar patterns with the model trained on e1minus enhancers, or the model trained on the whole data (Fig. 10). The features that are high ranking are still high ranking in both sets and predictive in the same directions. There are some differences in the transcription factors important between the two sets, but given the low accuracy of the e2plus data, we are not putting too much weight on the transcription factors that are only found in the e2plus model.

**Figure 10.**
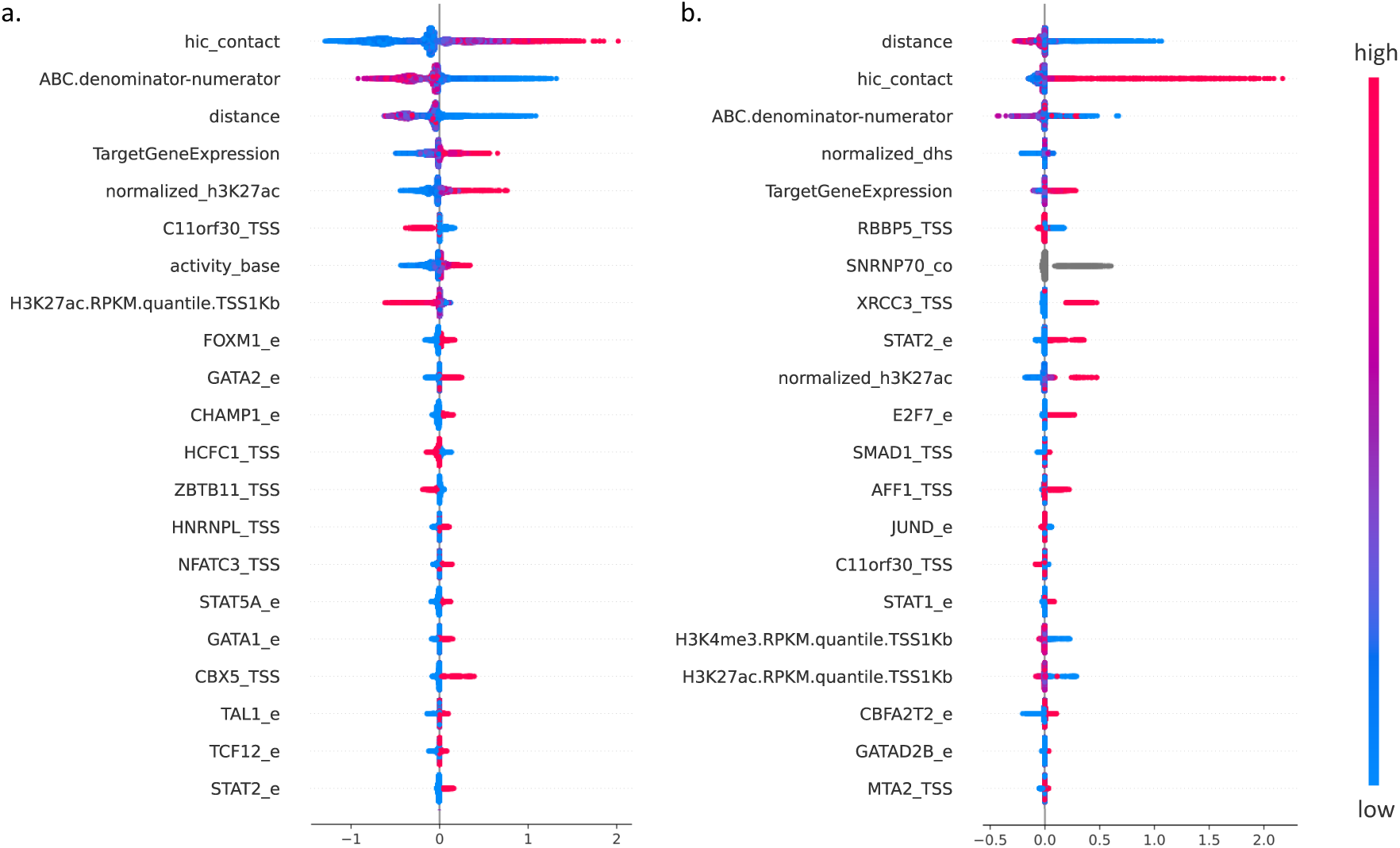
SHAP values for model trained on EPs with vs without direct Hi-C contact. SHAP values showing important feature contributions for model trained on **A)** EPs with direct contact (e1minus data) and **B)** EPs without direct contact (e2plus data).

We did notice an interesting difference in the feature rest-of-ABC (ABC denominator-numerator), that showed a distinct pattern between the e1minus and e2plus enhancers. Only in the e2plus enhancers, we saw a subset of enhancer-promoter pairs that showed opposite trend from the rest of the data, *i.e.* stronger positive prediction when the rest-of-ABC is larger (Fig. 11 B-C). We have more confidence in this difference, because, i) the pattern is replicated in the whole data across more data points (Fig. 11A), ii) the same subset of enhancer-promoter pairs show an interesting pattern going against the trend with the H3K4me3 at the TSS as well (Fig. 12). Even though generally the H3K4me3 mark at the TSS shows good correlation with the H3K27ac mark at the TSS, this particular pattern only showed up with H3K4me3, but not with H3K27ac (Fig. 12 vs. Fig. 6).

**Figure 11.**
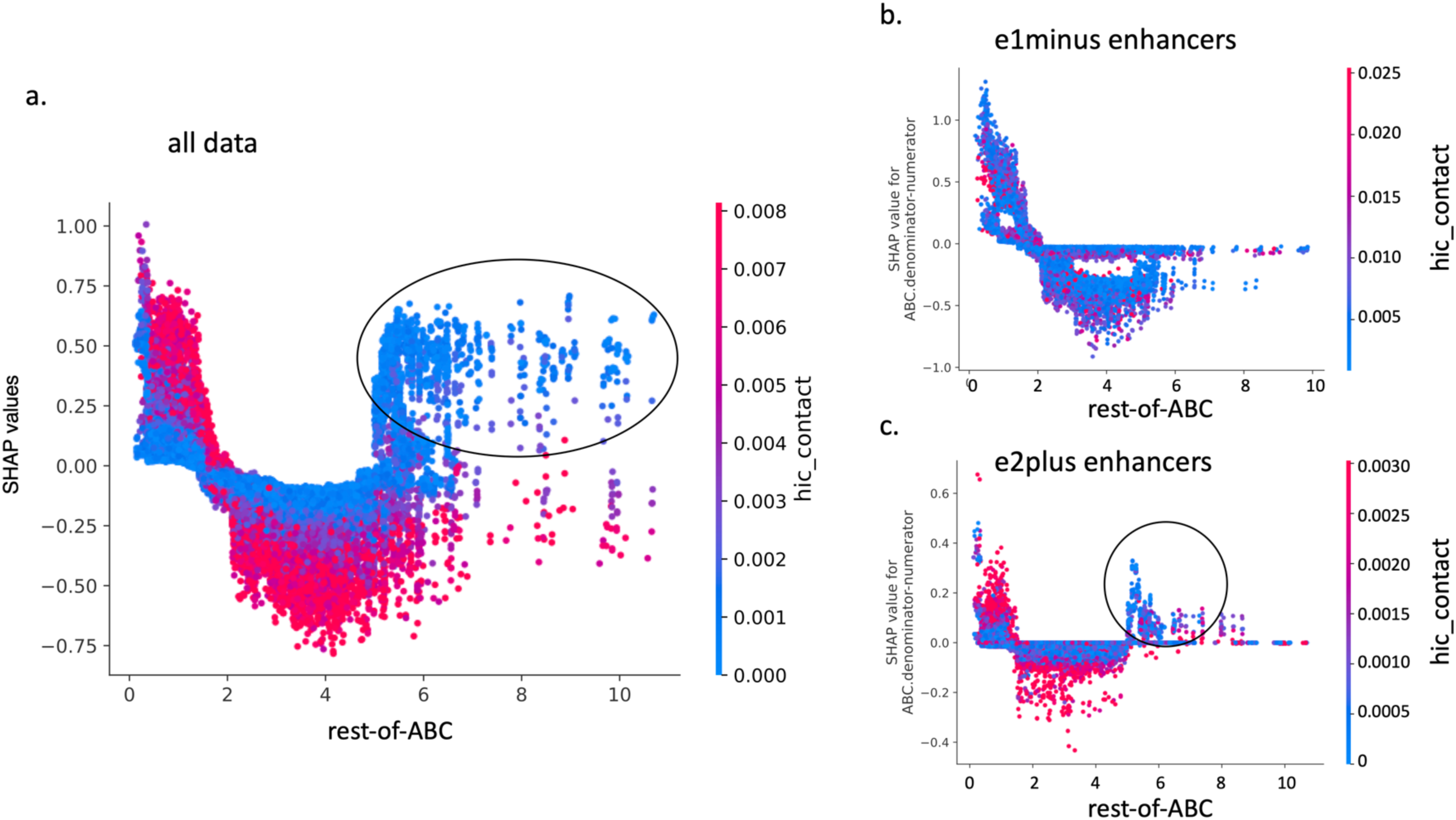
A distinct class of functional EP pairs revealed based on the feature rest-of-ABC. **A)** SHAP values for the feature rest-of-ABC (ABC denominator – numerator) of the model trained on the total dataset. For the rest of the data, large value of rest-of-ABC indicates strong or many enhancers nearby and predicts nonfunctional EP pairs, except for this subset of EPs highlighted with an oval. **B)** The model trained on EP pairs with direct contact (e1minus data) is missing this pattern. **C)** This pattern is recaptured in the model trained on EP pairs without direct contact (e2plus data).

**Figure 12.**
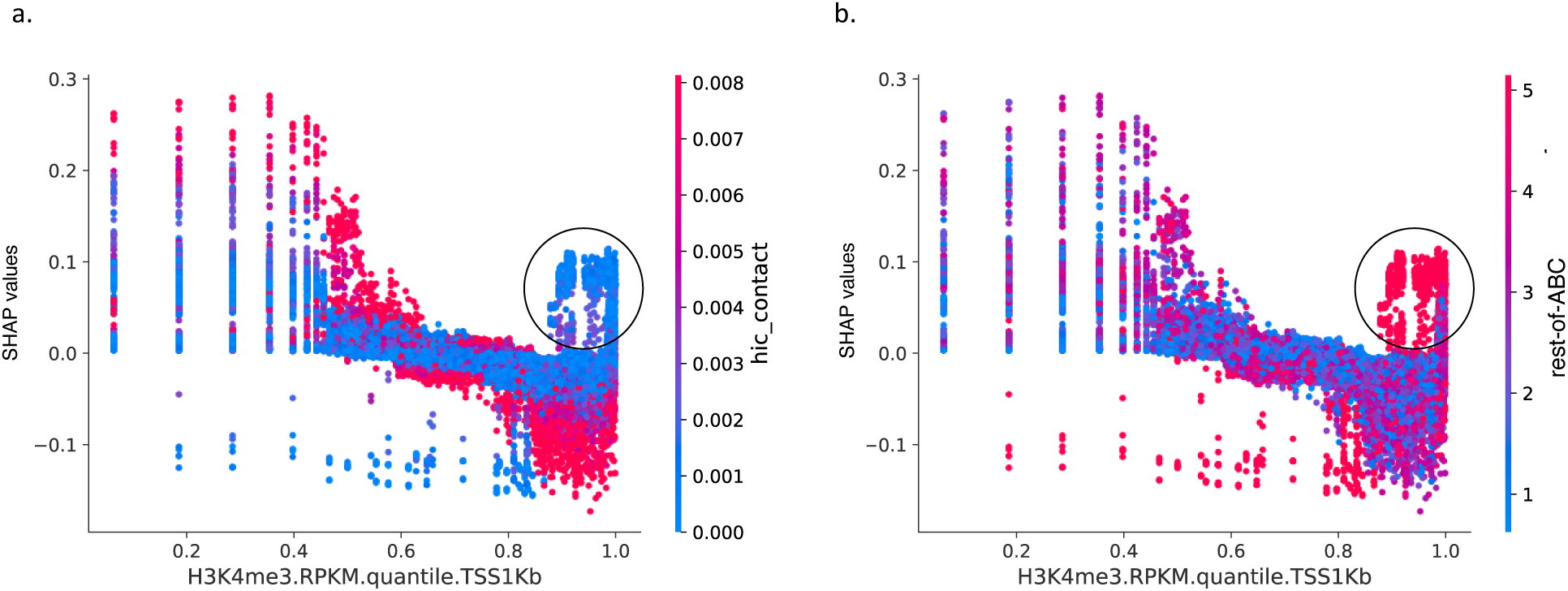
A distinct class of functional EP pairs and H3K4me3 at the promoter. The same class of indirect but functional EP pairs also show distinct pattern of SHAP values for H3K4me3 at the promoter. **A)** SHAP plot of promoter H3K4me3 marks with color scaled by Hi-C contact between enhancer and promoter and **B)** rest-of-ABC (ABC denominator - numerator). For the rest of the data, strong H3K4me3 at the promoter predicts nonfunctional EP pairs, except for this subset of EPs without direct contact and strong/many enhancers nearby.

What the model is predicting is that for a subset of enhancer-promoter pairs with weaker Hi-C contact, unlike the rest of the data, having other stronger enhancers nearby, and having a strong H3K4me3 peak at the TSS is contributing to the prediction of a functional EP pair. Based on these observations, we went back to our training data from Gasperini2019, to find the subset of positive cases that are under these conditions. We found 16 positive cases which satisfied all four conditions: i) Significant by Gasperini et al. ii) rest-of-ABC score (ABC denominator – numerator) > 5, iii) KR normalized Hi-C contact < 0.003, iv) quantile normalized H3K4me3 at the TSS > 0.8.

Table 3 lists those 16 cases. Of the 16, six were clustered enhancers targeting the same gene *ATG2A*, and four were clustered enhancers targeting genes in the Histone cluster 1, either *HIST1H4H* or *HIST1H2AC*. The histone genes and the gene *TBC1D17* were identified as part of the 704 genes that are regulated by super-enhancers in K562 (Hnisz et al., 2013). All other genes were also regulated by super-enhancers according to dbSuper, but not in the K562 cell line. For assurance, we checked our blocked Cross Validation design, and confirmed that all of the histone cluster 1 genes and enhancers on chromosome 6 from 25Mb to 29Mb were part of a single fold (fold 3) and not split across folds. The gene *ATG2A* is an interesting case, where there is a total of 22 enhancers tested by Gasperini et al., and six of those enhancers were determined to be positive. All six enhancers are included in Table 3, and are atypical in that they are not in direct contact with the gene’s promoter.

**Table 3.**
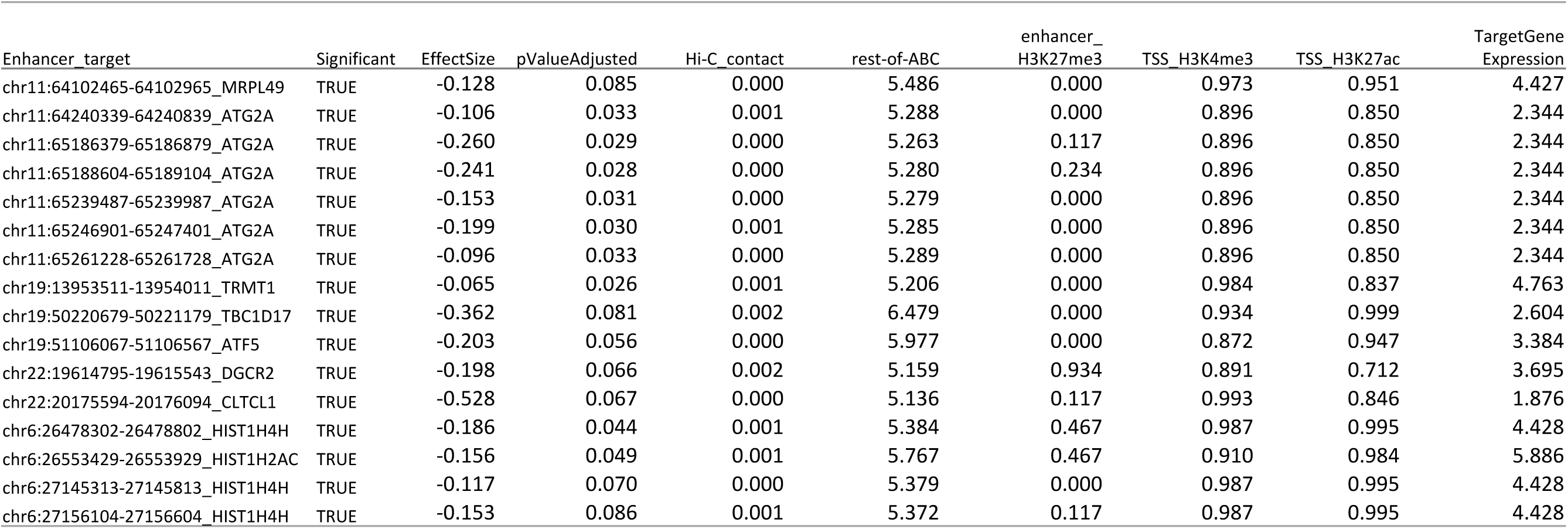
Examples of a distinct class of EP pairs. . Cases of significant EP pairs without direct contact that meet the following criteria: i) Significant by Gasperini et al. ii) rest-of-ABC (ABC score denominator – numerator) > 5, iii) KR normalized Hi-C contact < 0.003, iv) quantile normalized H3K4me3 at the TSS > 0.8.

## 4 Discussion

Distal enhancers and their role in transcriptional regulation have been studied for a long time, but each individual study has been limited to a combination of a handful of perturbation on specific enhancer loci. On the other hand, genomic studies have accumulated large amounts of biochemical assays that narrowed down the candidate regulatory elements, and gave meaning to the largely unknown areas of the genome, but they have been limited to clusterings and correlations and few had been functionally verified. But, with progress in technology, we have reached a point where now functional studies are being conducted at a genome-wide scale, and these parallel enhancer perturbation studies have accumulated individual enhancer perturbations and transcription read outs in the order of 10,000s of data points, that applying machine learning to the problem of transcriptional regulation specificity has now become practically feasible.

By integrating the numerous TF ChIP-seq data from ENCODE, with the experimental study by Gasperini et al. we find that we can train a model that is still limited in its accuracy, but nevertheless useful for gaining insight on how each feature contributes to the model outcome, and for generating hypotheses. The trained model corroborates the current model for enhancer-promoter prediction in many ways, highlighting the importance of interaction between the enhancer and promoter in physical space, and the importance of the activating chromatin marks such as H3K27ac at the enhancer. But, it also provides novel insights, showing that integrating further TF information improves the prediction, and pointing us towards the TFs that make a difference in this particular cell line/tissue context.

A few features were noteworthy. The observation that EMSY and HCFC1 was associated with negative prediction, but with higher activating chromatin marks at the TSS was unexpected. Understanding of chromatin modifiers enriched at the TSS is turning out to be more complicated, with many chromatin-associated proteins that were initially known to be either a repressor or an activator to reveal dual roles. EMSY was first characterized as a repressor (Hughes-Davies et al., 2003) that interacts with BRCA2, but subsequent reports on EMSY knock-outs and rescue experiments suggest EMSY can also activate transcription (Varier et al., 2016). Likewise, HCFC1 (HCF-1) is known to associate with both activator and repressor E2F proteins and to play different roles in different stages of the cell cycle regulation (Tyagi et al. MolCell 2007). In our model, the correlated features of EMSY, HCFC1 (HCF-1) ChIP-seq peaks and highly activating chromatin marks of H3K27ac and H3K4me3 at the TSS all lead to lower probability of predicting a functional enhancer for that target TSS. This is consistent with the recent report on the two classes of promoters by Bergman et al. that showed that promoters of housekeeping genes called P2s with activating motifs had decreased responsiveness to distal enhancers (Bergman et al., 2022). In fact, the ChIP-seq peaks of the two transcription factors HCFC1 and EMSY was identified in Bergman et al. among the enriched features that distinguished P2 promoters. We show that this pattern is replicated in a different experiment using CRISPR interference, compared to the massively parallel reporter assay that was used in Bergman et al. (Bergman et al., 2022). We found a few unexpected TFs that showed up as important in predicting positive enhancers, one of which was the CHAMP1 peak at the enhancer. This let us hypothesize that CHAMP1 is involved in protecting the chromosome integrity under replication transcription conflict. Enhancers are known to create R-loops similar to gene bodies, which can cause DNA damage by creating a barrier to replication fork progression. ChIP-seq peaks of CHAMP1 was observed to co-occur with peaks of FOXM1, which is active during late S phase, at the functional enhancers. The fact that a DNA repair gene is predictive of functional enhancers is at first puzzling. But it is consistent with the observation that functional enhancers generate higher level of transcripts and produce more R loops and may indicate that they are at higher risk for DNA damage. Although not part of the top 20 features (Fig. 4), the ChIP-seq peak of XRCC3, another gene with function in homologous recombinational repair, was also one of the important features based on its SHAP value. Its presence at the TSS was also positively predictive of functional enhancer-promoter pairs.

On the other hand, we found that the TFs that are well-known to bind to enhancers, and are routinely used to identify candidate enhancers, are curiously missing in our importance ranks. For example, we had included the ChIP-seq data for co-activators such as CREBBP (CBP), EP300 (P300), BRD4, and CEBPB, but none of them were determined to be important in predicting functional enhancers in this particular dataset. Another set of features deemed not important in our model was the co-binding of identical TFs between enhancers and promoters. We included such features, hypothesizing that TFs may mediate the chromatin interaction between enhancers and promoters, and if so, such TFs would show co-occurring peaks at the enhancers and promoters. But, co-occurring TF peaks were largely absent from the top important features, with SNRNP70 being the only exception. The co-occurring peaks of SNRNP70 showed up among the top 20 SHAP values. Recently, U1 snRNP was shown to tether and mobilize lncRNAs including enhancer RNAs to chromatin in a transcription-dependent manner (Yin et al., 2020).

Finally, although the majority of the enhancers were consistent with the overall patterns we described above, we found a distinct class of enhancers that went against such patterns. These were enhancers that did not exhibit a direct contact with the promoter measured by Hi-C, and instead were indirectly connected via a chain of enhancer to enhancer contacts. Opposite of the typical enhancers, these were more likely to have an effect when there were strong/numerous other enhancers around, and when the TSS had a strong activating mark of H3K4me3, but not necessarily of H3K27ac. Clusters of enhancers centered around the histone cluster 1 on chromosome 6, and the gene *ATG2A* are good examples of these atypical enhancers identified in the Gasperini experiment. One possible alternative explanation for these cases, other than our hypothesis of regulation through enhancer chain of contact, is that these enhancers are regulating another protein, that then regulates *ATG2A* through a trans effect. But, if that is the case, we don’t see why it would show association with strength of the other enhancers in the nearby region.

Our trained model is limited by its model system which is the K562 cell line. We observe that a lot of the positive data points in the Gasperini data are associated with cell cycle genes, including the histone gene cluster that is partially driven by the indirect enhancers described above. Since the transcriptional regulation of the histone cluster is somewhat specialized, it may not be an optimal model for studying regular enhancer driven regulation. In that sense *ATG2A* may be the better model to use for studying indirect enhancer regulation. Likewise, the DNA repair genes that are predictive of functional enhancers maybe a phenomenon that is only found in cancerous cells that are under high DNA replication stress. K562 cells are also aneuploids for most of the genome. These chromosomal aberrations are larger in scale than the Hi-C contacts we are measuring here, so it shouldn’t affect our results (chromosome 9 is excluded from our analysis due to the large-scale chromosomal inversion), but it may reduce the power of many of our genomic measurements, such as Hi-C, and ChIP-seq peaks, when homologous aneuploid chromosomes have underlying differences that are masked by aligning to a diploid genome.

A more central limitation would be that, since the increased performance relies on the TFs that are potentially cell type specific, the model would not achieve the same prediction accuracy on other cells or tissues. This is both the strength and weakness of the model, in that it becomes more accurate to the system, and loses generalizability. But, hopefully the knowledge that we learned from the model can be transferred to other systems.

## Acknowledgements

We would like to thank Corinne E. Sexton for her insights.

## Funding

This work was supported by the National Science Foundation under Grant No. 1750532.

### Conflict of Interest

none declared.

## Notes

### Competing Interest Statement

The authors have declared no competing interest.

## References

1. Akiba, T., Sano, S., Yanase, T., Ohta, T., & Koyama, M. (2019). Optuna: A next-generation hyperparameter optimization framework. 2623–2631.

2. Alexander, J. M., Guan, J., Li, B., Maliskova, L., Song, M., Shen, Y., Huang, B., Lomvardas, S., & Weiner, O. D. (2019). Live-cell imaging reveals enhancer-dependent Sox2 transcription in the absence of enhancer proximity. ELife, 8, e41769. https://doi.org/10.7554/eLife.41769

3. Bergman, D. T., Jones, T. R., Liu, V., Ray, J., Jagoda, E., Siraj, L., Kang, H. Y., Nasser, J., Kane, M., Rios, A., Nguyen, T. H., Grossman, S. R., Fulco, C. P., Lander, E. S., & Engreitz, J. M. (2022). Compatibility rules of human enhancer and promoter sequences. Nature, 607(7917), Article 7917. https://doi.org/10.1038/s41586-022-04877-w

4. Cao, F., Zhang, Y., Cai, Y., Animesh, S., Zhang, Y., Akincilar, S. C., Loh, Y. P., Li, X., Chng, W. J., Tergaonkar, V., Kwoh, C. K., & Fullwood, M. J. (2021). Chromatin interaction neural network (ChINN): A machine learning-based method for predicting chromatin interactions from DNA sequences. Genome Biology, 22(1), 226. https://doi.org/10.1186/s13059-021-02453-5

5. Cao, Q., Anyansi, C., Hu, X., Xu, L., Xiong, L., Tang, W., Mok, M. T. S., Cheng, C., Fan, X., Gerstein, M., Cheng, A. S. L., & Yip, K. Y. (2017). Reconstruction of enhancer–target networks in 935 samples of human primary cells, tissues and cell lines. Nature Genetics, 49(10), 1428–1436. https://doi.org/10.1038/ng.3950

6. Chen, T., & Guestrin, C. (2016). XGBoost: A Scalable Tree Boosting System. Proceedings of the 22nd ACM SIGKDD International Conference on Knowledge Discovery and Data Mining, 785–794. https://doi.org/10.1145/2939672.2939785

7. de Almeida, B. P., Reiter, F., Pagani, M., & Stark, A. (2022). DeepSTARR predicts enhancer activity from DNA sequence and enables the de novo design of synthetic enhancers. Nature Genetics, 54(5), 613–624. https://doi.org/10.1038/s41588-022-01048-5

8. Erwin, G. D., Oksenberg, N., Truty, R. M., Kostka, D., Murphy, K. K., Ahituv, N., Pollard, K. S., & Capra, J. A. (2014). Integrating Diverse Datasets Improves Developmental Enhancer Prediction. PLOS Computational Biology, 10(6), e1003677. https://doi.org/10.1371/journal.pcbi.1003677

9. Fudenberg, G., Kelley, D. R., & Pollard, K. S. (2020). Predicting 3D genome folding from DNA sequence with Akita. Nature Methods, 17(11), 1111–1117. https://doi.org/10.1038/s41592-020-0958-x

10. Fulco, C. P., Nasser, J., Jones, T. R., Munson, G., Bergman, D. T., Subramanian, V., Grossman, S. R., Anyoha, R., Patwardhan, T. A., Nguyen, T. H., Kane, M., Doughty, B., Perez, E. M., Durand, N. C., Stamenova, E. K., Aiden, E. L., Lander, E. S., & Engreitz, J. M. (2019). Activity-by-Contact model of enhancer specificity from thousands of CRISPR perturbations. BioRxiv, 529990. https://doi.org/10.1101/529990

11. Gasperini, M., Hill, A. J., McFaline-Figueroa, J. L., Martin, B., Kim, S., Zhang, M. D., Jackson, D., Leith, A., Schreiber, J., Noble, W. S., Trapnell, C., Ahituv, N., & Shendure, J. (2019). A Genome-wide Framework for Mapping Gene Regulation via Cellular Genetic Screens. Cell, 176(1), 377–390.e19. https://doi.org/10.1016/j.cell.2018.11.029

12. Hempel, M., Cremer, K., Ockeloen, C. W., Lichtenbelt, K. D., Herkert, J. C., Denecke, J., Haack, T. B., Zink, A. M., Becker, J., Wohlleber, E., Johannsen, J., Alhaddad, B., Pfundt, R., Fuchs, S., Wieczorek, D., Strom, T. M., van Gassen, K. L. I., Kleefstra, T., Kubisch, C., … Lessel, D. (2015). De Novo Mutations in CHAMP1 Cause Intellectual Disability with Severe Speech Impairment9. The American Journal of Human Genetics, 97(3), 493–500. https://doi.org/10.1016/j.ajhg.2015.08.003

13. Hnisz, D., Abraham, B. J., Lee, T. I., Lau, A., Saint-André, V., Sigova, A. A., Hoke, H. A., & Young, R. A. (2013). Super-Enhancers in the Control of Cell Identity and Disease. Cell, 155(4), 934–947. https://doi.org/10.1016/j.cell.2013.09.053

14. Hughes-Davies, L., Huntsman, D., Ruas, M., Fuks, F., Bye, J., Chin, S.-F., Milner, J., Brown, L. A., Hsu, F., Gilks, B., Nielsen, T., Schulzer, M., Chia, S., Ragaz, J., Cahn, A., Linger, L., Ozdag, H., Cattaneo, E., Jordanova, E. S., … Kouzarides, T. (2003). EMSY Links the BRCA2 Pathway to Sporadic Breast and Ovarian Cancer.Cell, 115(5), 523–535. https://doi.org/10.1016/S0092-8674(03)00930-9

15. Itoh, G., Kanno, S., Uchida, K. S. K., Chiba, S., Sugino, S., Watanabe, K., Mizuno, K., Yasui, A., Hirota, T., & Tanaka, K. (2011). CAMP (C13orf8, ZNF828) is a novel regulator of kinetochore–microtubule attachment. The EMBO Journal, 30(1), 130–144. https://doi.org/10.1038/emboj.2010.276

16. Kaul, A., Bhattacharyya, S., & Ay, F. (2020). Identifying statistically significant chromatin contacts from Hi-C data with FitHiC2. Nature Protocols, 15(3), 991–1012. https://doi.org/10.1038/s41596-019-0273-0

17. Kursa, M. B., & Rudnicki, W. R. (2010). Feature Selection with the Boruta Package. Journal of Statistical Software, 36(11), 1–13. https://doi.org/10.18637/jss.v036.i11

18. Li, F., Sarangi, P., Iyer, D. R., Feng, H., Moreau, L., Nguyen, H., Clairmont, C., & D’Andrea, A. D. (2022). CHAMP1 binds to REV7/FANCV and promotes homologous recombination repair. Cell Reports, 40(9). https://doi.org/10.1016/j.celrep.2022.111297

19. Lim, B., & Levine, M. S. (2021). Enhancer-promoter communication: Hubs or loops? Current Opinion in Genetics & Development, 67, 5–9. https://doi.org/10.1016/j.gde.2020.10.001

20. Lundberg, S. M., & Lee, S.-I. (2017). A unified approach to interpreting model predictions. Advances in Neural Information Processing Systems, 30.

21. Martinez-Ara, M., Comoglio, F., van Arensbergen, J., & van Steensel, B. (2022). Systematic analysis of intrinsic enhancer-promoter compatibility in the mouse genome. Molecular Cell, 82(13), 2519–2531.e6. https://doi.org/10.1016/j.molcel.2022.04.009

22. Miele, A., & Dekker, J. (2008). Long-range chromosomal interactions and gene regulation. Molecular BioSystems, 4(11), 1046–1057. https://doi.org/10.1039/b803580f

23. Palstra, R.-J., Tolhuis, B., Splinter, E., Nijmeijer, R., Grosveld, F., & de Laat, W. (2003). The β-globin nuclear compartment in development and erythroid differentiation. Nature Genetics, 35(2), Article 2. https://doi.org/10.1038/ng1244

24. Rowley, M. J., & Corces, V. G. (2018). Organizational principles of 3D genome architecture. Nature Reviews Genetics, 19(12), 789–800. https://doi.org/10.1038/s41576-018-0060-8

25. Sandhu, K. S., Li, G., Poh, H. M., Quek, Y. L. K., Sia, Y. Y., Peh, S. Q., Mulawadi, F. H., Lim, J., Sikic, M., Menghi, F., Thalamuthu, A., Sung, W. K., Ruan, X., Fullwood, M. J., Liu, E., Csermely, P., & Ruan, Y. (2012). Large-Scale Functional Organization of Long-Range Chromatin Interaction Networks. Cell Reports, 2(5), 1207–1219. https://doi.org/10.1016/j.celrep.2012.09.022

26. Schraivogel, D., Gschwind, A. R., Milbank, J. H., Leonce, D. R., Jakob, P., Mathur, L., Korbel, J. O., Merten, C. A., Velten, L., & Steinmetz, L. M. (2020). Targeted Perturb-seq enables genome-scale genetic screens in single cells. Nature Methods, 17(6), 629–635. https://doi.org/10.1038/s41592-020-0837-5

27. Schwessinger, R., Gosden, M., Downes, D., Brown, R. C., Oudelaar, A. M., Telenius, J., Teh, Y. W., Lunter, G., & Hughes, J. R. (2020). DeepC: predicting 3D genome folding using megabase-scale transfer learning. Nature Methods, 17(11), 1118–1124. https://doi.org/10.1038/s41592-020-0960-3

28. Song, W., Sharan, R., & Ovcharenko, I. (2019). The first enhancer in an enhancer chain safeguards subsequent enhancer-promoter contacts from a distance. Genome Biology, 20(1), 197. https://doi.org/10.1186/s13059-019-1808-y

29. Spilianakis, C. G., & Flavell, R. A. (2004). Long-range intrachromosomal interactions in the T helper type 2 cytokine locus. Nature Immunology, 5(10), Article 10. https://doi.org/10.1038/ni1115

30. Thibodeau, A., Márquez, E. J., Shin, D.-G., Vera-Licona, P., & Ucar, D. (2017). Chromatin interaction networks revealed unique connectivity patterns of broad H3K4me3 domains and super enhancers in 3D chromatin. Scientific Reports, 7(1), Article 1. https://doi.org/10.1038/s41598-017-14389-7

31. Tolhuis, B., Palstra, R.-J., Splinter, E., Grosveld, F., & de Laat, W. (2002). Looping and Interaction between Hypersensitive Sites in the Active β-globin Locus. Molecular Cell, 10(6), 1453–1465. https://doi.org/10.1016/S1097-2765(02)00781-5

32. Varier, R. A., de Santa Pau, E. C., van der Groep, P., Lindeboom, R. G. H., Matarese, F., Mensinga, A., Smits, A. H., Edupuganti, R. R., Baltissen, M. P., Jansen, P. W. T. C., ter Hoeve, N., van Weely, D. R., Poser, I., van Diest, P. J., Stunnenberg, H. G., & Vermeulen, M. (2016). Recruitment of the Mammalian Histone-modifying EMSY Complex to Target Genes Is Regulated by ZNF131 *. Journal of Biological Chemistry, 291(14), 7313–7324. https://doi.org/10.1074/jbc.M115.701227

33. Whalen, S., Schreiber, J., Noble, W. S., & Pollard, K. S. (2022). Navigating the pitfalls of applying machine learning in genomics. Nature Reviews Genetics, 23(3), 169–181. https://doi.org/10.1038/s41576-021-00434-9

34. Whalen, S., Truty, R. M., & Pollard, K. S. (2016). Enhancer–promoter interactions are encoded by complex genomic signatures on looping chromatin. Nature Genetics, 48(5), 488–496. https://doi.org/10.1038/ng.3539

35. Xi, W., & Beer, M. A. (2018). Local epigenomic state cannot discriminate interacting and non-interacting enhancer– promoter pairs with high accuracy. PLOS Computational Biology, 14(12), e1006625. https://doi.org/10.1371/journal.pcbi.1006625

36. Xiao, J. Y., Hafner, A., & Boettiger, A. N. (2021). How subtle changes in 3D structure can create large changes in transcription. ELife, 10, e64320. https://doi.org/10.7554/eLife.64320

37. Yang, B., Liu, F., Ren, C., Ouyang, Z., Xie, Z., Bo, X., & Shu, W. (2017). BiRen: Predicting enhancers with a deep-learning-based model using the DNA sequence alone. Bioinformatics, 33(13), 1930–1936. https://doi.org/10.1093/bioinformatics/btx105

38. Yin, Y., Lu, J. Y., Zhang, X., Shao, W., Xu, Y., Li, P., Hong, Y., Cui, L., Shan, G., Tian, B., Zhang, Q. C., & Shen, X. (2020). U1 snRNP regulates chromatin retention of noncoding RNAs. Nature, 580(7801), 147–150. https://doi.org/10.1038/s41586-020-2105-3

